# Molecular analysis of prostate cancer: prostate tissue and urine proteomics based approach

**DOI:** 10.1101/2021.01.02.423577

**Authors:** Amrita Mitra, Rajdeep Das, Surya Kant Choubey, Amit Kumar Mandal

**Affiliations:** Clinical Proteomics Unit, Division of Molecular Medicine, St. John’s Research Institute, St. John’s National Academy of Health Sciences, 100ft Road, Koramangala, Bangalore – 560034; Department of Urology and Renal Transplantation, St. Johns Medical College & Hospital, St. John’s National Academy of Health Sciences, 100ft Road, Koramangala, Bangalore – 560034

## Abstract

Benign prostatic hyperplasia (BPH) and prostate cancer are the most frequently diagnosed conditions in men above 60 years. BPH manifests as benign enlargement of the prostate gland causing pain and difficulty in micturition, often associated with other lower urinary tract symptoms. On the other hand, prostate cancer might initially set in as a tumor of no clinical significance, but can ultimately develop into an aggressively metastatic cancer at later stages. Due to overlapping symptomatic manifestations with BPH, it might be difficult to diagnose prostate cancer and differentiate it from BPH. Screening for prostate cancer is based on examining the levels of prostate specific antigen (PSA), a clinical biomarker for prostate cancer in blood. However, several reported cases indicate that PSA might lack the sensitivity and specificity required to differentiate between the cancerous and benign conditions of prostate. Therefore, in the absence of non-invasive biomarkers in a diagnostic setup, an intensely invasive surgical removal of a part of the tissue and subsequent histopathologial examination is the only available procedure to confirm the cancer. In this study, we used tissue and urine proteomics platforms to profile respective proteomes in both clinical conditions. We observed that latent transforming growth factor binding protein was significantly under-expressed in the both tissue and urine proteome of prostate cancer. We propose that the down-regulation of the latent transforming growth factor binding protein 4 might be explored in a large set of patients with prostate cancer to develop a non-invasive urine based biomarker for the diagnosis of prostate cancer.

## Introduction

Prostate cancer is the one of the commonly diagnosed malignancies worldwide and is the second most common type of cancer in men over the age of 60 years [1, 2]. The disease initially sets in as a slow growing tumor, barely associated with any clinical significance, but might ultimately develop into an aggressively metastatic cancer in the later stages [3]. The major ambiguity in the screening of prostate cancer lies in its overlapping symptomatic manifestations with another age-related prostatic condition, benign prostatic hyperplasia (BPH). Both these conditions are associated with clinical symptoms such as abdominal pain, difficult in micturition, loss of blood in urine and other associated lower urinary tract symptoms (LUTS) [4]. The routine primary screening of prostate cancer involves digital rectal examination (DRE) and transrectal ultrasonography (TRUS). However, both DRE and TRUS might not be very sensitive methods to diagnose distantly located tumors in the prostate gland [5, 6]. In addition, for the routine diagnosis of prostate cancer, prostate specific antigen (PSA), a blood biomarker, is quantified in blood samples to diagnose the presence of cancer [7]. The normal level of PSA in blood is 4 ng/mL. An increase in the level of PSA from the normal is clinically associated with the presence of cancer. However, PSA screening has been reported to lack sensitivity and specificity for the disease. Firstly, a rise in the levels of PSA was observed to be associated with non-malignant disorders such as BPH, infection or chronic inflammation of the prostate [8]. Secondly, studies have also reported the prevalence of prostate cancer in patients with normal PSA levels [9]. The lack of high specificity and sensitivity of the available diagnostic tools necessitates the discovery of a more reliable screening procedure that can efficiently distinguish between the cancerous and benign conditions of the prostate gland.

Based on the differential expression of proteins in clinical samples, mass spectrometry based proteomics approach is an ideal platform to investigate for biomarkers of a disease. Additionally, various post-translation modifications of proteins that are commonly observed in different disease states, can also be analysed by this method. Urine proteomics based approach is an appropriate strategy to develop a biomarker for prostate cancer where the sample can be collected non-invasively. Till date, some studies have reported individual set of biomarkers in tissues and urine samples respectively for prostate cancer [10–14]. However, it is important to correlate the expression of the marker protein in both tissue and urine proteomes simultaneously such that the prostate cancer biomarker can be unambiguously used in the diagnosis of prostate cancer in future.

In the present study, we used mass spectrometry based proteomics platform for profiling the expression of proteins in tissues and urine samples collected from patients with prostate cancer and compared them with the tissue and urine proteomes of the benign condition (BPH). Subsequently, the proteomic alterations in prostate cancer were correlated between the tissue and urine proteomes to observe the changes that commonly occurred in both the tissue and urine proteomes in prostate cancer.

## Materials and methods

### Materials

Analytical grade ethanol was obtained from Merck (Germany). Sequencing grade bovine trypsin was purchased from Sigma Aldrich (St. Louis, MO). Water, acetonitrile and TFA were of LC−MS grade and obtained from Honeywell (California, USA). All other chemicals used were of analytical grade.

### Ethics statement

The study was approved by the Institutional Ethical Committee (IEC approval no. 27/2014), St. John’s Medical College, Bangalore, Karnataka, India.

### Selection of subjects

Patients presenting with prostate cancer and benign prostatic hyperplasia to the Department of Urology at St. John’s Medical College Hospital, Bangalore, were invited to participate in the study. Written consents were obtained from patients prior to collection of samples.

### Collection and processing of tissue samples from patients with BPH and prostate cancer

Twenty prostate needle core biopsy samples and a prostate tissue bit were collected in normal saline from ten patients suspected with prostate cancer. Out of the collected needle core biopsies, seven were diagnosed with prostate cancer using routine histopathological examination. Additionally, twenty tissue bits from ten patients with BPH undergoing the procedure of transurethral resection of the prostate (TURP) were also collected. One core/bit of prostate tissue samples was lyophilized for 4 hours. The lyophilized tissue was crushed to powder. 0.2 mg of the crushed sample was suspended in tissue lysis buffer containing 10 mM tris HCl, 0.15 M NaCl, 1 mM EDTA in phosphate buffered saline, pH 7.4 with 1% β-octyl glucopyranoside. 10 μL of protease inhibitor cocktail was added to the suspension and subsequently sonicated for 30 min with intermittent cooling. After sonication, the solution was incubated for 1 hour at 4 °C with mild agitation. The suspension was centrifuged at 14,700 × g for 45 min and the supernatant was dialyzed overnight with frequent changes against 50 mM NH_4_HCO_3_ buffer, pH 7.4. After dialysis, the protein concentration was measured by Bradford assay method according to the manufacturer’s protocol and followed by in-solution digestion.

### Collection and processing of urine samples from patients with BPH and prostate cancer

Samples were collected from a patient with BPH to as a control and seven samples were collected from patients with prostate cancer. From each subject 80-100 ml of urine was collected into a sterile container containing protease inhibitors cocktail and sodium azide to prevent proteolysis and bacterial growth. The urine samples were centrifuged at 1575 × g for 15 mins at room temperature to remove the cell debris and particulate matter. The proteins present in the supernatant were precipitated by the addition of ethanol at 4 °C using urine to solvent ratio of 1:9 and incubated for 8 hours. The protein precipitate was collected by centrifugation at 18,514 × g for 15 mins at 4 °C. The obtained pellet was air dried and dissolved into a buffer containing 7 M urea, 2 M thiourea. The protein solution was then dialyzed overnight with frequent changes against 50 mM NH_4_HCO_3_ buffer, pH 7.4. After dialysis, the protein concentration of the solution was assessed by Biorad assay method following the manufacturer’s protocol.

### In-solution digestion of the proteins from tissue and urine samples

For digestion, 25 μg of protein was used. Samples were denatured by using 0.2% Rapigest at 80 °C for 15 min. Subsequently, the disulphide linkages of proteins were reduced by 5 mM dithiothreitol for 30 min at 60 °C, followed by alkylation of the reduced -SH groups with 10 mM iodoacetamide for 30 min in dark at room temperature. Modified sequencing grade trypsin, reconstituted in 50 mM NH_4_HCO_3_, pH 7.4, was added to the protein extracts at an enzyme:substrate ratio of 1:10 and incubated overnight at 37 °C. Following this, the samples were acidified in 0.5% (v/v) formic acid (FA) and incubated at 37 °C for 90 mins to facilitate hydrolysis of Rapigest detergent. Samples were then be centrifuged for 45 min at 4000 × g and the supernatant was collected and diluted to obtain a final amount of 0.2 μg/μL of the digested protein. The samples were spiked with 50 femtomoles/μl of internal standard yeast enolase (WATERS, UK). Subsequently, the samples were carried forward for nanoLC-ESI-MS analysis.

### nanoLC fractionation

The samples were analyzed at Proteomics Facility, Centre for Cellular and Molecular Platforms (CCAMP), National Centre for Biological Sciences (NCBS), Bangalore. 3 μL of the sample containing a total of 600 ng of the digested proteins and 150 femtomoles of enolase were fractionated in EASY-nanoLC 1200 UHPLC system (Thermo Scientific, Germany), calibrated using Pierce™ LTQ ESI Positive Ion Calibration Solution (Thermo Scientific, Germany). Briefly, 600 ng of total tryptic digest of the experimental sample was loaded directly into a C18 Acclaim™ PepMap™ Rapid Separation Liquid Chromatography column (3 μm, 100 Å × 75 μm × 50 cm). The aqueous phase consisted of water/0.1% FA (solvent A) and the organic phase comprised of acetonitrile/0.1% FA (solvent B). The flow rate of solvents through the column was 300 nL/min. A 90 min run program was designed, starting with a linear gradient of solvent B from 5-40% for 70 min, followed by 40-80% of solvent B for 8 min, and finally 80-95% of solvent B for 12 min. The column temperature was maintained at 40 °C during the run in all experiments. The eluted peptides were analyzed in a mass spectrometer with an orbitrap mass analyser (Orbitrap Fusion™ mass spectrometer, Thermo Scientific, San Jose, California, USA).

### Data acquisition

The data were acquired in the mass range of 375−1700 m/z at 120000 FWHM resolution (at 200 m/z). The instrument was operated in positive ion mode with a source temperature of 350 °C, capillary voltage of 2.0 kV. Poly-siloxane (445.12003 m/z), used as lock mass, was continuously infused throughout the acquisition. The instrument was set to run in top speed mode with 3 s cycles for the survey and the MS/MS scans. After a survey scan, tandem MS was performed on the most abundant precursors exhibiting charge states from 2 to 7 with the intensity greater than 1.0e^4^, by isolating them in the quadrupole. High Collision Dissociation (HCD) fragmentation was applied with 35% collision energy and resulting fragments were detected using the rapid scan rate in the ion trap. The automatic gain control (AGC) target for MS/MS was set to 4.0e^5^ and the maximum injection time limited to 50 msec. The experimental samples were run in duplicate.

### Proteomics Data Analysis

The acquired data was processed using Thermo Scientific™ Proteome Discoverer™ software version 2.1. MS/MS spectra were searched with the SEQUEST^®^ HT engine against human proteome database incorporated with yeast enolase sequence details. During analysis, the parameters used were as follows: proteolytic enzyme, trypsin; missed cleavages allowed, two; minimum peptide length, 6 amino acids. Carbamidomethylation (+57.021 Da) at cysteine residues was kept as a fixed modification. Methionine oxidation (+15.9949 Da) and protein N-terminal acetylation (+42.0106 Da) were set as variable modifications. The mass tolerances for precursor and product ions were kept at 10 ppm and 0.6 Da respectively. Peptide spectral matches (PSM) were validated using the Percolator^®^ algorithm, based on q-values at a 1% FDR. Peptides were filtered based on the high FDR values. Protein groups were further identified with at least two unique peptides.

The software calculated the peak area of any given protein as the average of the peak areas of three most abundant distinct peptides identified for the protein. The peptides must have different sequences to be considered distinct. Peptides with different charge states or modification variants of the same sequence are considered the same peptide [15]. Several proteins were characterized and identified in the experiment with the values of their corresponding peak areas. From the values obtained for the peak areas of the proteins, their relative quantities in femtomoles were calculated based on the internal standard yeast enolase, whose average peak area corresponded to 150 femtomoles, the total amount injected in the column. Following this, the data was normalized by dividing the amount of each protein by the summation of the amounts of all proteins present in the sample. Subsequently, the proteins which appeared in both the technical duplicates of each sample were filtered out. From this list, a further filtration of proteins was performed to take forward only those proteins that appeared in at least five out of the six biological replicates of patients with prostate cancer. The averages of the amounts of each protein between the duplicate runs were calculated and the standard deviation was noted. Coefficient of variation between the duplicate runs of the samples was estimated using the following formula:

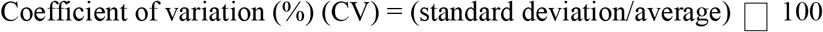

The proteins were again filtered to keep only those that appeared with a CV of less than equal to 15% between the duplicate runs of the samples. In addition, among the biological replicates, a broader degree of coefficient of variation of less than equal to 30% was selected and the proteins passing the criteria were selected for further analysis. From the list, the amounts of each protein in the biological replicates were averaged and were compared for the fold difference in expression. Proteins that displayed with greater than equal two fold change in expression between the BPH (control) and prostate cancer (experimental sample) sets, as an acceptable measure of significant differential expression in a disease condition against control, were reported [16].

## Results and discussion

Prostate cancer is one of the most frequently diagnosed cancer and a leading cause of mortality among males in developed countries [17]. However, due to its overlapping symptomatic manifestations with BPH, it might be difficult to diagnose cancer in the prostate and differentiate it from the benign condition [18, 19]. Therefore, in the absence of non-invasive biomarkers in the diagnostic setup, a surgical removal of a part of the tissue and subsequent histopathologial examination of it might be the only available procedure to confirm cancer. Although serum PSA evaluation is used to detect prostate cancer, it has been reported with lack of sensitivity and specificity to differentiate between the cancerous and benign conditions of the prostate in several cases [20, 21]. In the present study, the tissue and urine proteomics of prostate cancer in comparison to that of BPH showed differential expression levels of a protein, latent transforming growth factor binding protein 4, an activator of transforming growth factor β1 (TGF-β1) known to act as a tumor suppressor in early stages of cancers. as We observed that the protein was significantly underexpressed by 0.2 folds. We propose that the latent transforming growth factor binding protein 4 might be a potential candidate to explore as a non-invasive biomarker for the diagnosis of prostate cancer in the clinical setup.

### Tissue Proteomics

Tissue proteomics performed with human prostate tissues diagnosed with either cancer or BPH revealed that forty four proteins were significantly overexpressed and thirty eight proteins were underexpressed in the prostate cancer tissue proteome compared to that with the benign condition (Tables S1 and S2). A pie chart pictorial representation of these proteins plotted by using Panther Classification System software and classified based of their localization in the tissues has been provided in figures 1A and 1B.

**Figure 1:**
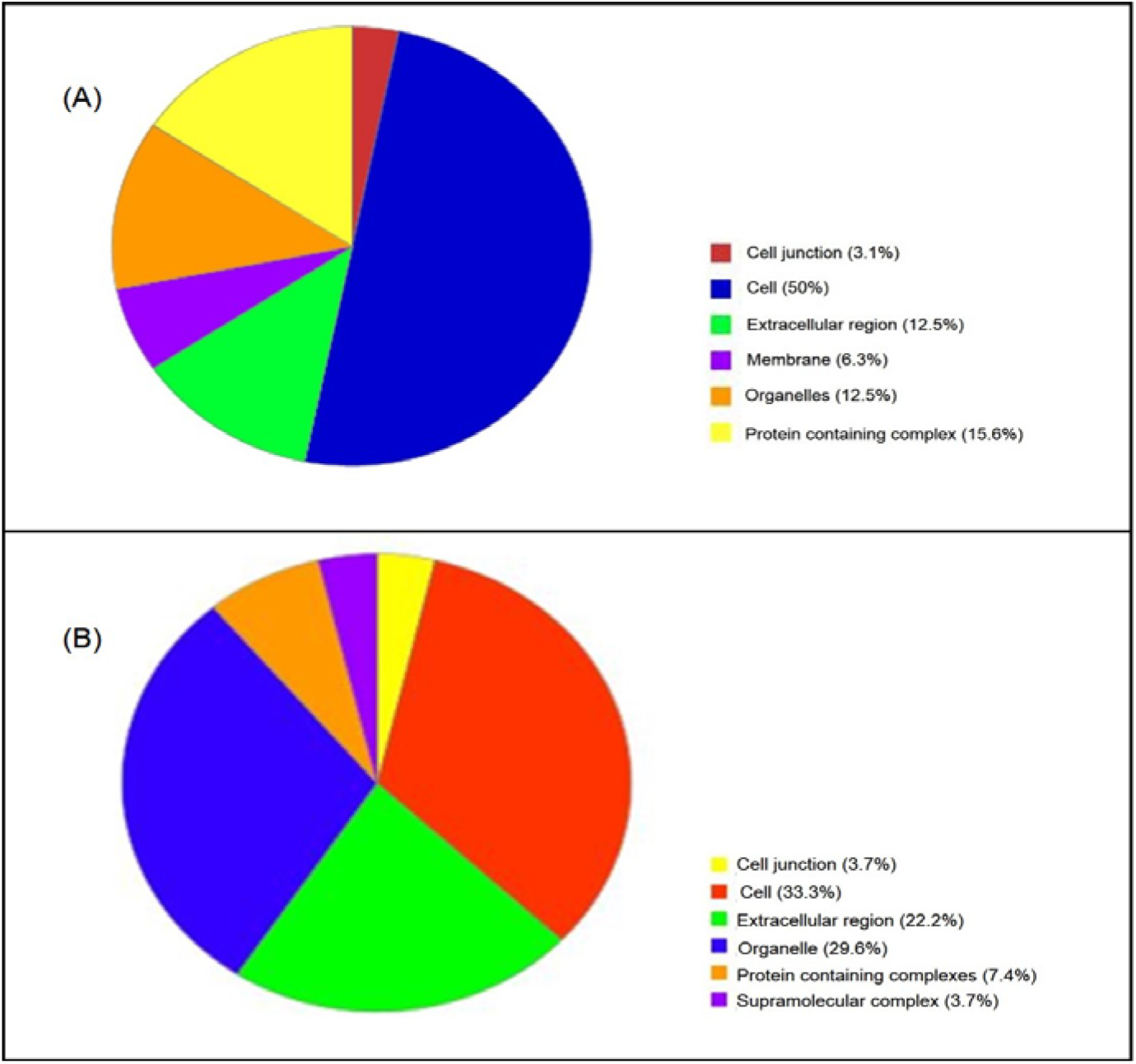
Pie chart representing the proteins with expressional alteration in prostate cancer tissue proteome compared to BPH and classified on the basis of their localization in the tissues. Panels A and B show the localization of the overexpressed and underexpressed proteins in prostate cancer tissue proteome respectively

Additionally, figures 2A and 2B shows the pie diagram of the overexpressed and underexpressed proteins respectively represented on the basis of their involvement in different biological processes.

**Figure 2:**
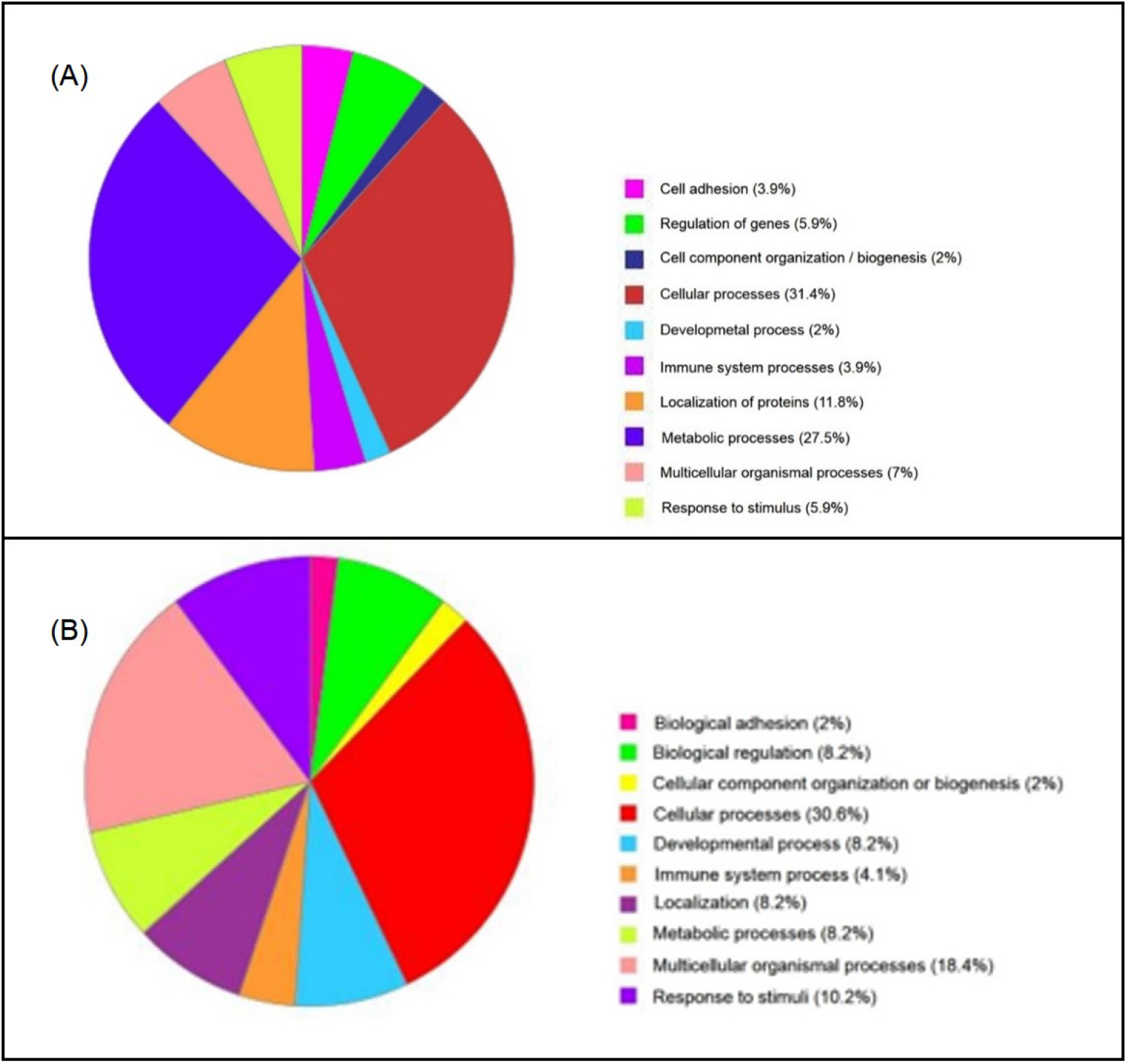
Pie chart representing the proteins with expressional alteration in prostate cancer tissue proteome compared to BPH and classified on the basis of their involvement in biological processes in the tissues. Panels A and B show the biological processes of the overexpressed and underexpressed proteins in prostate cancer tissue proteome respectively

Upon analysis of figures 2A and 2B, we observed that a major number of the proteins that were altered in the prostate cancer tissue proteome were involved in various cellular processes. The overexpressed proteins were observed to be majorly involved in protein metabolism, protein modification and protein folding processes (Figure 3A), whereas the underexpressed proteins were involved in the organization of different cellular components like chromatin, organelles, protein subunits, etc. Another major class of underexpressed proteins were observed to be associated management of cellular stress and cell signal transduction processes involving G-protein coupled receptor pathways, apoptotic signalling pathways, etc (Figure 3B).

**Figure 3:**
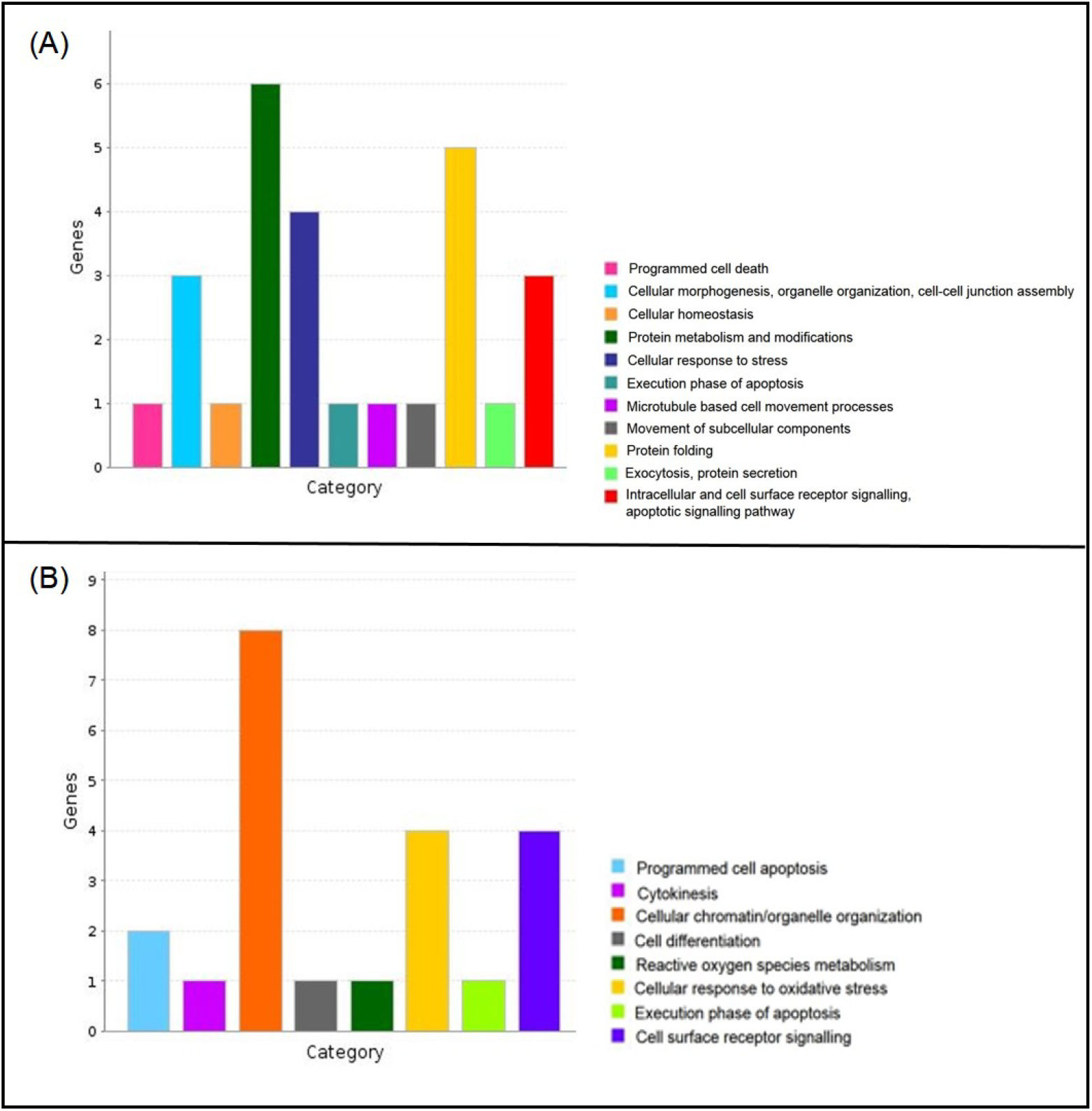
Sub-categorization of the various types of cellular processes of the genes of differentially expressed tissue proteins in prostate cancer. Panels A and B show the different cellular processes performed by a majority of the overexpressed and underexpressed proteins in prostate cancer tissue proteome respectively

### Urine Proteomics

Urine proteomics was performed to identify, characterize and quantify the proteins that underwent a differential expression in the urine proteome in patients with prostate cancer when compared to that in patients with BPH. We observed that seven proteins showed a significant increase in expression in the urine proteome of prostate cancer in comparison with BPH, whereas fifteen proteins represented with a significant reduction in expression (Tables S3 and S4). Figures 4A and 4B show the groups of proteins with altered expression in prostate cancer in a pie chart representation based on their biological functions.

**Figure 4:**
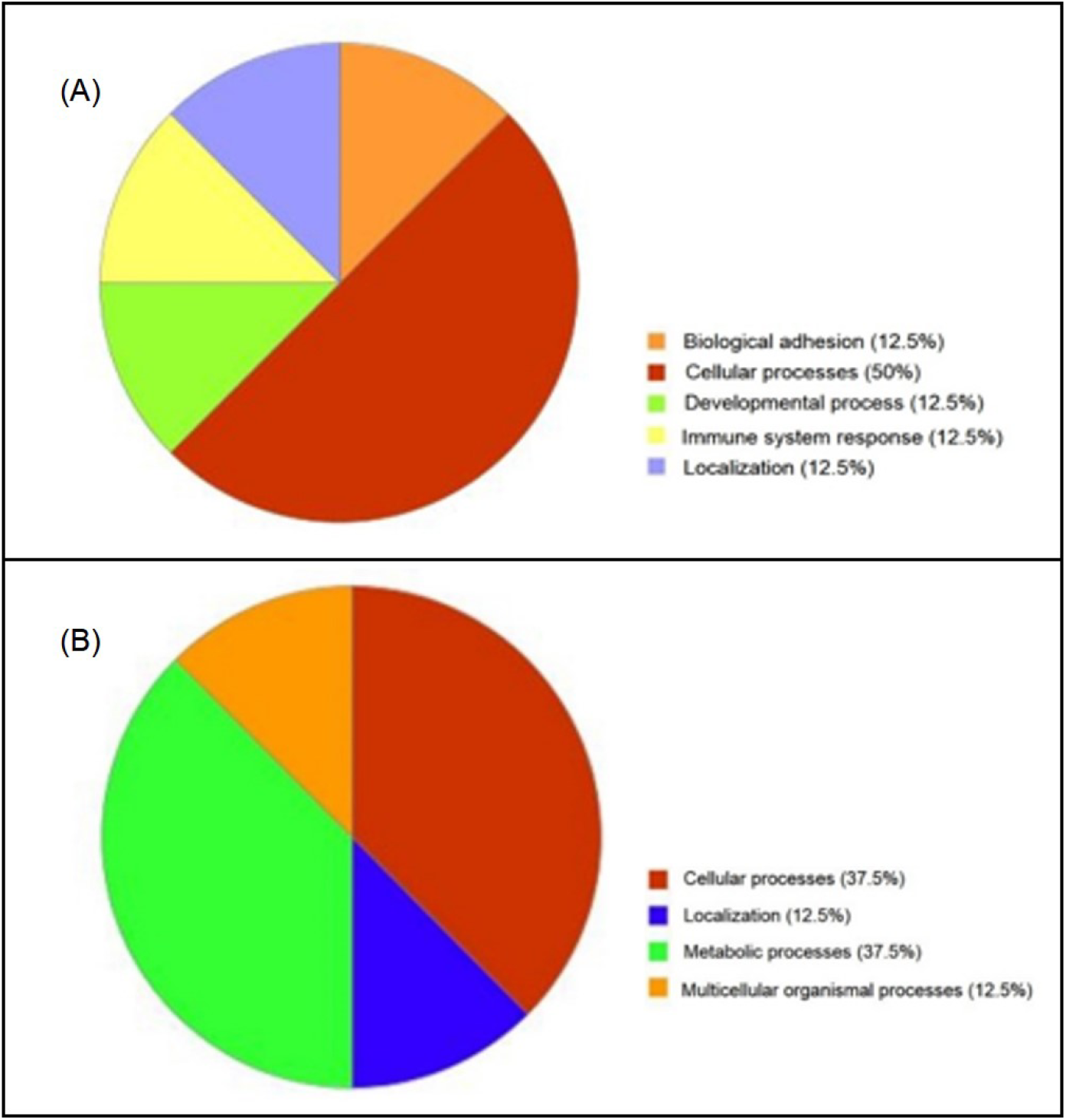
Pie chart representing the proteins with expressional alteration in prostate cancer urine proteome compared to BPH and classified on the basis of their involvement in biological processes in the prostate gland. Panels A and B show the biological processes of the overexpressed and underexpressed proteins in prostate cancer urine proteome respectively

A major number of overexpressed proteins were observed to be involved in various intracellular functions, whereas smaller fractions of proteins were observed to be involved in enhancement of cell adhesion processes, immune system related processes like leukocyte differentiation, localization of several transmembrane proteins, etc. We further analysed the class of proteins related to cellular processes and observed that a major part of the proteins were associated with cellular signal transduction, majorly via the RAS mediated protein signalling pathway, a signal transduction pathway that controls cell proliferation, migration and survival [22]. The other proteins were observed to be associated with organization of cellular components like organelles and autophagosomes, along with assembling subunits of different proteins (Figure 5A). Upon analysis of the underexpresssed proteins of the prostate cancer urine proteome, we observed that majority of the proteins were involved in various cellular and metabolic processes. A smaller fraction of these proteins were associated with biological functions such as the processes associated with sensory perception in the nervous system. Another small fraction of proteins were associated with localization of macromolecules like lipids to the cell membranes. We further analysed the larger fractions of the proteins associated with metabolic and cellular processes and observed that the proteins associated with cellular processes were involved in the metabolism of cell membrane lipids, that is, glycolipids and sphingolipids. On the other hand, the class of proteins associated with metabolic processes were observed to be involved in pathways majorly of carbohydrate and lipid metabolism (Figure 5B).

**Figure 5:**
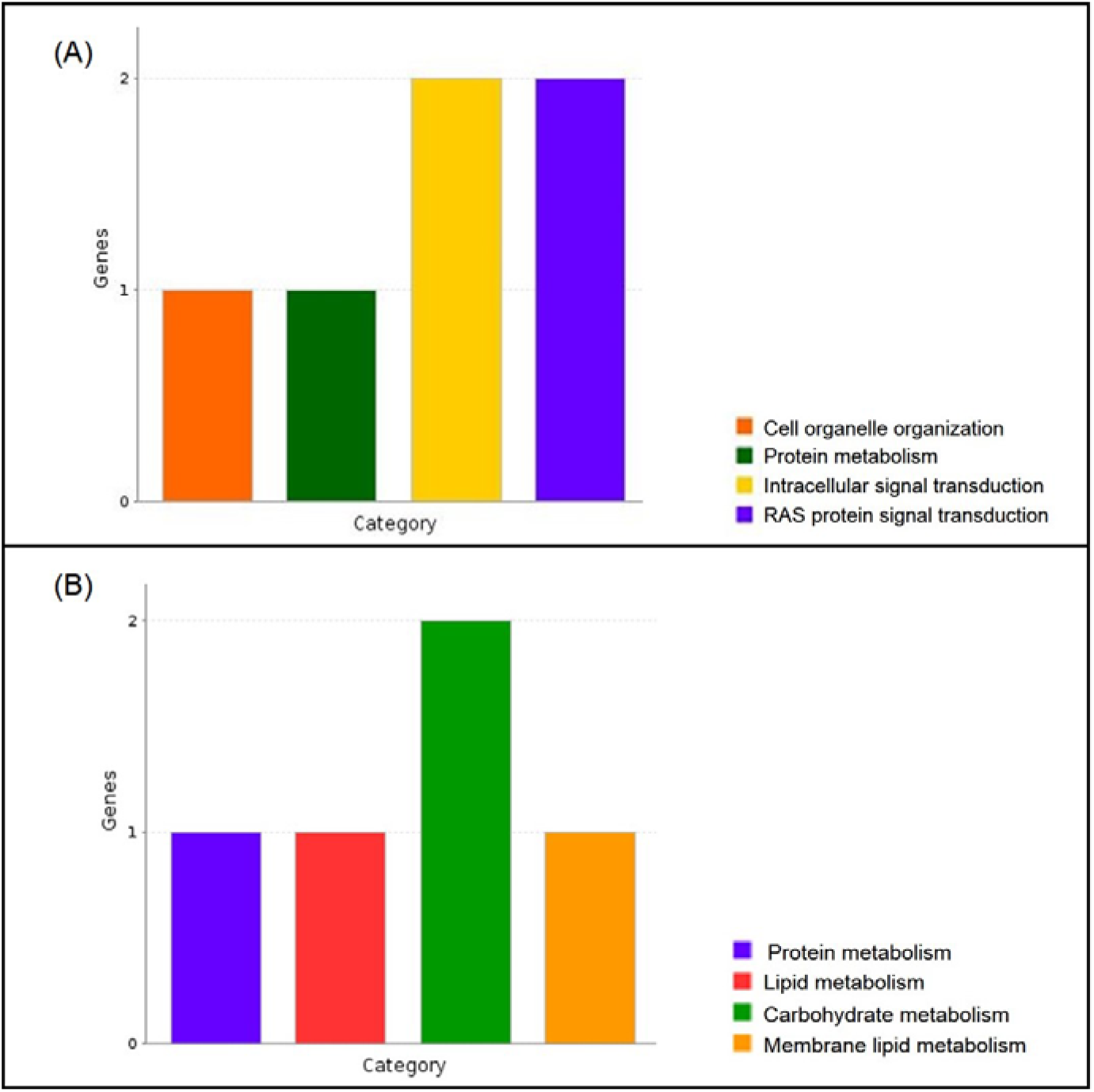
Sub-categorization of the various types of cellular and metabolic processes of the genes of differentially expressed tissue proteins in prostate cancer urine proteome. Panel A shows the different cellular processes performed by a majority of the overexpressed urinary proteins and panel B shows the involvement of a major number of underexpressed urinary proteins in various metabolic processes in prostate cancer

### Comparison between tissue and urine proteomics

The differential expression analysis in the tissue and urine proteomes of patients with prostate cancer showed that a protein, latent-transforming growth factor beta-binding protein 4, coded by the gene LTBP-4, was significantly underexpressed in both the tissue and urine proteomes of these patients (0.2 folds). It has been reported that latent transforming growth factor β-binding protein is an extracellular matrix glycoprotein associated molecule and is involved in the correct folding and secretion of transforming growth factor β1 (TGF-β1), a pluripotent cytokine associated with cellular proliferation, differentiation, apoptosis, etc. It has also been reported that TGF-βs, in general, potently inhibit cellular proliferation and regulate cellular differentiation and adhesion. TGF-βs are also associated with ECM production and degradation. At early stages of cancer development, TGF-βs act as tumor suppressors. Therefore, downregulation of LTBP-4 gene, involved in the activation of TGF-β1, promotes cancer development and progression [23]. It has also been well documented that LTBP-4 gene is downregulated in cancers of specific tissues including esophagus [24], colorectal cancer [25] and mammary carcinomas [26]. Downregulation of the LTBP-4 gene is also associated with lung dysfunction [23], cardiomyopathy [25] and muscular dystrophy [27]. However, the underexpression of the latent-transforming growth factor beta-binding protein 4 has not been reported in prostate cancer. In our study, we observed for the first time the underexpression of the protein, latent-transforming growth factor beta-binding protein 4, in the tissue and urine proteomes in prostate cancer. This extends the tissue specificity of the differential expression of latent-transforming growth factor beta-binding protein 4 to the cancer of the human prostate gland.

## Conclusions

In the present study, we observed the global changes that occur in the tissue and urine proteomes of patients with prostate cancer compared to the benign enlargement of the prostate, BPH. The group of proteins enlisted as significantly altered in the prostate cancer tissue and urine proteomes serves to unambiguously distinguish between the two conditions of the prostate gland with similar clinical manifestations, that is, prostate cancer and BPH. Additionally, the significant underexpression of latent-transforming growth factor beta-binding protein 4 in both tissue and urine proteomes of patients with prostate cancer might be extremely important to explore in a clinical setup in future in order to develop this protein as a specific and sensitive urinary biomarker for prostate cancer which can be examined in patients in an absolutely non-invasive manner.

## Supporting information

Supplementary doc

## Acknowledgements

We acknowledge all the volunteers for providing tissue samples for the study. We acknowledge Indian Council of Medical Research (ICMR), Govt. of India for funding the urine proteomics study and Rajiv Gandhi University of Health Sciences (RGUHS), Bangalore, for funding the tissue proteomics study.

## References

1. Taitt, H.E.: Global trends and prostate cancer: A review of incidence, detection, and mortality as influenced by race, ethnicity, and geographic location. American Journal of Men’s Health. 12, 1807–1823 (2018).

2. Bray, F., Ferlay, J., Soerjomataram, I., Siegel, R.L., Torre, L.A., Jemal, A.: Global cancer statistics 2018: GLOBOCAN estimates of incidence and mortality worldwide for 36 cancers in 185 countries. A Cancer Journal for Clinicians. 68, 394–424 (2018).

3. Hoimes, C.J., Kelly, W.K.: Redefining hormone resistance in prostate cancer. Therapeutic Advances in Medical Oncology. 2, 107–123 (2010).

4. Bhindi, A., Bhindi, B., Kulkarni, G.S., Hamilton, R.J., Toi, A., van der Kwast, T.H., Evans, A., Zlotta, A.R., Finelli, A., Fleshner, N.E.: Modern-day prostate cancer is not meaningfully associated with lower urinary tract symptoms: Analysis of a propensity score-matched cohort. Journal of the Canadian Urological Association. 11, 41–46 (2017).

5. Carvalhal, G.F., Smith, D.S., Mager, D.E., Ramos, C., Catalona, W.J.: Digital rectal examination for detecting prostate cancer at prostate specific antigen levels of 4 ng./ml. or less. The Journal of Urology. 161, 835–839 (1999).

6. Carter, H.B., Hamper, U.M., Sheth, S., Sanders, R.C., Epstein, J.I., Walsh, P.C.: Evaluation of transrectal ultrasound in the early detection of prostate cancer. The Journal of Urology. 142, 1008–1010 (1989).

7. Botchorishvili, G., Matikainen, M.P., Lilja, H.: Early prostate-specific antigen changes and the diagnosis and prognosis of prostate cancer. Current Opinion in Urology. 19, 221–226 (2009).

8. Stamey, T.A., Yang, N., Hay, A.R., McNeal, J.E., Freiha, F.S., Redwine, E.: Prostate-specific antigen as a serum marker for adenocarcinoma of the prostate. New England Journal of Medicine. 317, 909–916 (1987).

9. Morgentaler, A., Rhoden, E.L.: Prevalence of prostate cancer among hypogonadal men with prostate-specific antigen levels of 4.0 ng/mL or less. Urology. 68, 1263–1267 (2006).

10. Kristiansen, G.: Markers of clinical utility in the differential diagnosis and prognosis of prostate cancer. Modern Pathology. 31, 143–155 (2018).

11. Fiorentino, M., Capizzi, E. art, Loda, M.: Blood and tissue biomarkers in prostate cancer: State of the art. Urologic Clinics of North America. 37, 131–141 (2010).

12. Clinton, T.N., Bagrodia, A., Lotan, Y., Margulis, V., Raj, G. V., Woldu, S.L.: Tissue-based biomarkers in prostate cancer. Expert Review of Precision Medicine and Drug Development. 2, 249–260 (2017).

13. Fujita, K., Nonomura, N.: Urinary biomarkers of prostate cancer. International Journal of Urology. 25, 770–779 (2018).

14. Wu, D., Ni, J., Beretov, J., Cozzi, P., Willcox, M., Wasinger, V., Walsh, B., Graham, P., Li, Y.: Urinary biomarkers in prostate cancer detection and monitoring progression. Critical Reviews in Oncology/Hematology. 118, 15–26 (2017).

15. Silva, J.C., Gorenstein, M. V., Li, G.-Z., Vissers, J.P.C., Geromanos, S.J.: Absolute quantification of proteins by LCMSE: a virtue of parallel MS acquisition. Molecular & Cellular Proteomics. 5, 144–156 (2006).

16. Cho, H., Kim, H., Na, D., Kim, S.Y., Jo, D., Lee, D.: Meta-analysis method for discovering reliable biomarkers by integrating statistical and biological approaches: An application to liver toxicity. Biochemical and Biophysical Research Communications. 471, 274–281 (2016).

17. Rawla, P.: Epidemiology of prostate cancer. World Journal of Oncology. 10, 63–89 (2019).

18. Dai, X., Fang, X., Ma, Y., Xianyu, J.: Benign prostatic hyperplasia and the risk of prostate cancer and bladder cancer. Medicine. 95, e3493 (2016).

19. Miah, S., Catto, J.: BPH and prostate cancer risk. Indian Journal of Urology. 30, 214–218 (2014).

20. Fridriksson, J., Gunseus, K., Stattin, P.: Information on pros and cons of prostate-specific antigen testing to men prior to blood draw: A study from the National Prostate Cancer Register (NPCR) of Sweden. Scandinavian Journal of Urology and Nephrology. 46, 326–331 (2012).

21. Catalona, W.J., Loeb, S.: Prostate cancer screening and determining the appropriate prostate-specific antigen cutoff values. JNCCN Journal of the National Comprehensive Cancer Network. 8, 265–270 (2010).

22. Khosravi-Far, R., Der, C.J.: The Ras signal transduction pathway. Cancer and Metastasis Reviews. 13, 67–89 (1994).

23. Koli, K., Wempe, F., Sterner-Kock, A., Kantola, A., Komor, M., Hofmann, W.-K., von Melchner, H., Keski-Oja, J.: Disruption of LTBP-4 function reduces TGF-β activation and enhances BMP-4 signaling in the lung. The Journal of Cell Biology. 167, 123–133 (2004).

24. Bultmann, I., Conradi, A., Kretschmer, C., Sterner-Kock, A.: Latent transforming growth factor β-binding protein 4 is downregulated in esophageal cancer via promoter methylation. PLoS ONE. 8, e65614 (2013).

25. Sterner-Kock, A., Thorey, I.S., Koli, K., Wempe, F., Otte, J., Bangsow, T., Kuhlmeier, K., Kirchner, T., Jin, S., Keski-Oja, J., von Melchner, H.: Disruption of the gene encoding the latent transforming growth factor-beta binding protein 4 (LTBP-4) causes abnormal lung development, cardiomyopathy, and colorectal cancer. Genes & Development. 16, 2264–2273 (2002).

26. Kretschmer, C., Conradi, A., Kemmner, W., Sterner-Kock, A.: Latent transforming growth factor binding protein 4 (LTBP4) is downregulated in mouse and human DCIS and mammary carcinomas. Cellular Oncology. 34, 419–434 (2011).

27. Su, C.T., Huang, J.W., Chiang, C.K., Lawrence, E.C., Levine, K.L., Dabovic, B., Jung, C., Davis, E.C., Madan-Khetarpal, S., Urban, Z.: Latent transforming growth factor binding protein 4 regulates transforming growth factor beta receptor stability. Human Molecular Genetics. 24, 4024–4036 (2015).

