## Supplementary doc for "Molecular analysis of prostate cancer: prostate tissue and urine proteomics based approach"

Dr. Amit Kumar Mandal

Clinical Proteomics Unit

Division of Molecular Medicine

St. John's Research Institute,

St. John's National Academy of Health Sciences

100ft Road, Koramangala,

Bangalore – 560034, India

Present address:

Dr. Amit Kumar Mandal

Department of Biological Sciences,

Indian Institute of Science Education and Research Kolkata

Mohanpur, Nadia, West Bengal

India

Pin-741246

**Table S1:** List of proteins observed to be overexpressed using tissue proteomics by at least two folds in prostate cancer tissues compared to BPH tissues

| **Proteins** | **Accession No.** | **Gene code** | **Control (normalized and averaged)** | **Sample (normalized and averaged)** | **Fold change in expression (Sample/Control)** |
| --- | --- | --- | --- | --- | --- |
| Keratin, type I cytoskeletal 17 OS=Homo sapiens OX=9606 GN=KRT17 PE=1 SV=2 | Q04695 | KRT17 | 0.0001 | 0.0017 | 16.85 |
| 40S ribosomal protein S7 OS=Homo sapiens OX=9606 GN=RPS7 PE=1 SV=1 | P62081 | RPS7 | 0.0002 | 0.0019 | 8.30 |
| Succinate dehydrogenase [ubiquinone] flavoprotein subunit, mitochondrial OS=Homo sapiens OX=9606 GN=SDHA PE=1 SV=2 | P31040 | SDHA | 0.0008 | 0.0001 | 7.84 |
| Serpin H1 OS=Homo sapiens OX=9606 GN=SERPINH1 PE=1 SV=2 | P50454 | SERPINH1 | 0.0002 | 0.0014 | 7.75 |
| PDZ and LIM domain protein 5 OS=Homo sapiens OX=9606 GN=PDLIM5 PE=1 SV=5 | Q96HC4 | PDLIM5 | 0.0001 | 0.0007 | 6.48 |
| Cytochrome c oxidase subunit 2 OS=Homo sapiens OX=9606 GN=MT-CO2 PE=1 SV=1 | P00403 | MT-CO2 | 0.0001 | 0.0006 | 6.08 |
| 60 kDa heat shock protein, mitochondrial OS=Homo sapiens OX=9606 GN=HSPD1 PE=1 SV=2 | P10809 | HSPD1 | 0.0003 | 0.0018 | 6.00 |
| 2,4-dienoyl-CoA reductase, mitochondrial OS=Homo sapiens OX=9606 GN=DECR1 PE=1 SV=1 | Q16698 | DECR1 | 0.0002 | 0.0011 | 5.39 |
| Translocon-associated protein subunit delta OS=Homo sapiens OX=9606 GN=SSR4 PE=1 SV=1 | P43307 | SSR1 | 0.0001 | 0.0007 | 5.37 |
| Tryptase alpha/beta-1 OS=Homo sapiens OX=9606 GN=TPSAB1 PE=1 SV=1 | Q15661 | TPSAB1 | 0.0003 | 0.0017 | 4.94 |
| Delta(3,5)-Delta(2,4)-dienoyl-CoA isomerase, mitochondrial OS=Homo sapiens OX=9606 GN=ECH1 PE=1 SV=2 | Q13011 | ECH1 | 0.0002 | 0.0008 | 4.87 |
| Protein disulfide-isomerase OS=Homo sapiens OX=9606 GN=P4HB PE=1 SV=3 | P07237 | P4HB | 0.0017 | 0.0073 | 4.22 |
| Cytochrome b5 OS=Homo sapiens OX=9606 GN=CYB5A PE=1 SV=2 | P00167 | CYB5A | 0.0002 | 0.0007 | 3.99 |
| Isoform Sap-mu-9 of Prosaposin OS=Homo sapiens OX=9606 GN=PSAP | P07602 | PSAP | 0.0009 | 0.0032 | 3.61 |
| Adipocyte plasma membrane-associated protein OS=Homo sapiens OX=9606 GN=APMAP PE=1 SV=2 | Q9HDC9 | APMAP | 0.0001 | 0.0005 | 3.52 |
| Protein disulfide-isomerase A4 OS=Homo sapiens OX=9606 GN=PDIA4 PE=1 SV=2 | P13667 | PDIA4 | 0.0005 | 0.0018 | 3.47 |
| Dihydrolipoyllysine-residue succinyltransferase component of 2-oxoglutarate dehydrogenase complex, mitochondrial OS=Homo sapiens OX=9606 GN=DLST PE=1 SV=4 | P36957 | DLST | 0.0002 | 0.0008 | 3.28 |
| Enoyl-CoA hydratase, mitochondrial OS=Homo sapiens OX=9606 GN=ECHS1 PE=1 SV=4 | P30084 | ECHS1 | 0.0001 | 0.0003 | 3.14 |
| Myeloid-derived growth factor OS=Homo sapiens OX=9606 GN=MYDGF PE=1 SV=1 | Q969H8 | MYDGF | 0.0002 | 0.0007 | 3.06 |
| Isoform 2 of Vesicle-associated membrane protein-associated protein A OS=Homo sapiens OX=9606 GN=VAPA | Q9P0L0 | VAPA | 0.0001 | 0.0003 | 2.97 |
| Voltage-dependent anion-selective channel protein 1 OS=Homo sapiens OX=9606 GN=VDAC1 PE=1 SV=2 | P21796 | VDAC1 | 0.0002 | 0.0007 | 2.96 |
| Isoform 2B of Cytoplasmic dynein 1 intermediate chain 2 OS=Homo sapiens OX=9606 GN=DYNC1I2 | Q13409 | DYNC1I2 | 0.0001 | 0.0003 | 2.95 |
| Thioredoxin domain-containing protein 5 OS=Homo sapiens OX=9606 GN=TXNDC5 PE=1 SV=2 | Q8NBS9 | TXNDC5 | 0.0005 | 0.0014 | 2.92 |
| Cadherin-1 OS=Homo sapiens OX=9606 GN=CDH1 PE=1 SV=3 | P12830 | CDH1 | 0.0001 | 0.0003 | 2.92 |
| Electron transfer flavoprotein subunit alpha, mitochondrial OS=Homo sapiens OX=9606 GN=ETFA PE=1 SV=1 | P13804 | ETFA | 0.0001 | 0.0004 | 2.92 |
| Peroxiredoxin-4 OS=Homo sapiens OX=9606 GN=PRDX4 PE=1 SV=1 | Q13162 | PRDX4 | 0.0001 | 0.0004 | 2.87 |
| Peptidyl-prolyl cis-trans isomerase B OS=Homo sapiens OX=9606 GN=PPIB PE=1 SV=2 | P23284 | PPIB | 0.0010 | 0.0028 | 2.85 |
| Fibrinogen alpha chain OS=Homo sapiens OX=9606 GN=FGA PE=1 SV=2 | P02671 | FGA | 0.0007 | 0.0019 | 2.80 |
| Isoform 2 of Protein disulfide-isomerase A6 OS=Homo sapiens OX=9606 GN=PDIA6 | Q15084 | PDIA6 | 0.0008 | 0.0022 | 2.77 |
| 60S acidic ribosomal protein P2 OS=Homo sapiens OX=9606 GN=RPLP2 PE=1 SV=1 | P05387 | RPLP2 | 0.0006 | 0.0015 | 2.62 |
| Isoform 2 of Keratin, type II cytoskeletal 8 OS=Homo sapiens OX=9606 GN=KRT8 | P05787 | KRT8 | 0.0022 | 0.0056 | 2.59 |
| T-complex protein 1 subunit gamma OS=Homo sapiens OX=9606 GN=CCT3 PE=1 SV=4 | P49368 | CCT3 | 0.0002 | 0.0006 | 2.54 |
| Fructose-bisphosphate aldolase C OS=Homo sapiens OX=9606 GN=ALDOC PE=1 SV=2 | P09972 | ALDOC | 0.0002 | 0.0004 | 2.45 |
| Monocyte differentiation antigen CD14 OS=Homo sapiens OX=9606 GN=CD14 PE=1 SV=2 | P08571 | CD14 | 0.0001 | 0.0001 | 2.36 |
| 3-hydroxyacyl-CoA dehydrogenase type-2 OS=Homo sapiens OX=9606 GN=HSD17B10 PE=1 SV=3 | Q99714 | HSD17B10 | 0.0001 | 0.0002 | 2.32 |
| Decorin OS=Homo sapiens OX=9606 GN=DCN PE=1 SV=1 | P07585 | DCN | 0.0051 | 0.0117 | 2.28 |
| Isoform 2 of Calnexin OS=Homo sapiens OX=9606 GN=CANX | P27824 | CANX | 0.0006 | 0.0014 | 2.23 |
| Heterogeneous nuclear ribonucleoprotein F OS=Homo sapiens OX=9606 GN=HNRNPF PE=1 SV=3 | P52597 | HNRNPF | 0.0002 | 0.0004 | 2.22 |
| T-complex protein 1 subunit beta OS=Homo sapiens OX=9606 GN=CCT2 PE=1 SV=4 | P78371 | CCT2 | 0.0002 | 0.0005 | 2.21 |
| Dolichyl-diphosphooligosaccharide--protein glycosyltransferase subunit 1 OS=Homo sapiens OX=9606 GN=RPN1 PE=1 SV=1 | P04843 | RPN1 | 0.0002 | 0.0005 | 2.10 |
| Cytochrome c oxidase subunit 5A, mitochondrial OS=Homo sapiens OX=9606 GN=COX5A PE=1 SV=2 | P20674 | COX5A | 0.0002 | 0.0003 | 2.10 |
| Endoplasmin OS=Homo sapiens OX=9606 GN=HSP90B1 PE=1 SV=1 | P14625 | HSP90B1 | 0.0016 | 0.0032 | 2.08 |
| Fibrinogen gamma chain OS=Homo sapiens OX=9606 GN=FGG PE=1 SV=3 | P02679 | FGG | 0.0013 | 0.0028 | 2.07 |
| Serotransferrin OS=Homo sapiens OX=9606 GN=TF PE=1 SV=3 | P02787 | TF | 0.0015 | 0.0031 | 2.05 |

**Table S2:** List of proteins observed to be underexpressed using tissue proteomics by at least two folds in prostate cancer tissues compared to BPH tissues

| **Proteins** | **Accession No.** | **Gene code** | **Control (normalized and averaged)** | **Sample (normalized and averaged)** | **Fold change in expression (Sample/Control)** |
| --- | --- | --- | --- | --- | --- |
| Membrane primary amine oxidase OS=Homo sapiens OX=9606 GN=AOC3 PE=1 SV=3 | Q16853 | AOC3 | 0.0016 | 0.0008 | 0.49 |
| Annexin A5 OS=Homo sapiens OX=9606 GN=ANXA5 PE=1 SV=2 | P08758 | ANXA5 | 0.0026 | 0.0013 | 0.48 |
| Isoform 2 of Fibulin-2 OS=Homo sapiens OX=9606 GN=FBLN2 | P98095 | FBLN2 | 0.0002 | 0.0001 | 0.48 |
| Mimecan OS=Homo sapiens OX=9606 GN=OGN PE=1 SV=1 | P20774 | OGN | 0.0020 | 0.0009 | 0.46 |
| Superoxide dismutase [Cu-Zn] OS=Homo sapiens OX=9606 GN=SOD1 PE=1 SV=2 | P00441 | SOD1 | 0.0039 | 0.0016 | 0.42 |
| Epoxide hydrolase 1 OS=Homo sapiens OX=9606 GN=EPHX1 PE=1 SV=1 | P07099 | EPHX1 | 0.0004 | 0.0002 | 0.41 |
| Nidogen-2 OS=Homo sapiens OX=9606 GN=NID2 PE=1 SV=3 | Q14112 | NID2 | 0.0005 | 0.0002 | 0.41 |
| Collagen alpha-1(I) chain OS=Homo sapiens OX=9606 GN=COL1A1 PE=1 SV=5 | P02452 | CO1A1 | 0.0049 | 0.0020 | 0.40 |
| Isoform 2 of Tropomyosin alpha-3 chain OS=Homo sapiens OX=9606 GN=TPM3 | P06753 | TPM3 | 0.0029 | 0.0012 | 0.39 |
| EGF-containing fibulin-like extracellular matrix protein 1 OS=Homo sapiens OX=9606 GN=EFEMP1 PE=1 SV=2 | Q12805 | EFEMP1 | 0.0005 | 0.0002 | 0.38 |
| Guanine nucleotide-binding protein G(I)/G(S)/G(T) subunit beta-1 OS=Homo sapiens OX=9606 GN=GNB1 PE=1 SV=3 | P62873 | GNB1 | 0.0003 | 0.0001 | 0.38 |
| Sodium/potassium-transporting ATPase subunit alpha-1 OS=Homo sapiens OX=9606 GN=ATP1A1 PE=1 SV=1 | P05023 | ATP1A1 | 0.0003 | 0.0001 | 0.36 |
| Myosin regulatory light polypeptide 9 OS=Homo sapiens OX=9606 GN=MYL9 PE=1 SV=4 | P24844 | MYL9 | 0.0061 | 0.0022 | 0.36 |
| Cysteine and glycine-rich protein 1 OS=Homo sapiens OX=9606 GN=CSRP1 PE=1 SV=3 | P21291 | CSRP1 | 0.0083 | 0.0030 | 0.35 |
| Myosin regulatory light chain 12B OS=Homo sapiens OX=9606 GN=MYL12B PE=1 SV=2 | O14950 | MYL12B | 0.0008 | 0.0003 | 0.35 |
| Isoform 3 of Agrin OS=Homo sapiens OX=9606 GN=AGRN | O00468 | AGRN | 0.0001 | 0.0000 | 0.33 |
| Galectin-1 OS=Homo sapiens OX=9606 GN=LGALS1 PE=1 SV=2 | P09382 | LGALS1 | 0.0034 | 0.0011 | 0.33 |
| 4-trimethylaminobutyraldehyde dehydrogenase OS=Homo sapiens OX=9606 GN=ALDH9A1 PE=1 SV=3 | P49189 | ALDH9A1 | 0.0005 | 0.0002 | 0.33 |
| Annexin A6 OS=Homo sapiens OX=9606 GN=ANXA6 PE=1 SV=3 | P08133 | ANXA6 | 0.0017 | 0.0006 | 0.33 |
| Isoform LAMP-2C of Lysosome-associated membrane glycoprotein 2 OS=Homo sapiens OX=9606 GN=LAMP2 | P13473 | LAMP2 | 0.0008 | 0.0003 | 0.32 |
| Calponin-1 OS=Homo sapiens OX=9606 GN=CNN1 PE=1 SV=2 | P51911 | CNN1 | 0.0127 | 0.0039 | 0.31 |
| Caldesmon OS=Homo sapiens OX=9606 GN=CALD1 PE=1 SV=3 | Q05682 | CALD1 | 0.0008 | 0.0003 | 0.30 |
| Sarcolemmal membrane-associated protein OS=Homo sapiens OX=9606 GN=SLMAP PE=1 SV=1 | Q14BN4 | SLMAP | 0.0004 | 0.0001 | 0.29 |
| Core histone macro-H2A.1 OS=Homo sapiens OX=9606 GN=H2AFY PE=1 SV=4 | O75367 | H2AFY | 0.0007 | 0.0002 | 0.28 |
| Basement membrane-specific heparan sulfate proteoglycan core protein OS=Homo sapiens OX=9606 | P98160 | HSPG2 | 0.0011 | 0.0003 | 0.27 |
| Myosin light polypeptide 6 OS=Homo sapiens OX=9606 GN=MYL6 PE=1 SV=2 | P60660 | MYL6 | 0.0156 | 0.0039 | 0.25 |
| Isoform 4 of Four and a half LIM domains protein 1 OS=Homo sapiens OX=9606 GN=FHL1 | Q13642 | FHL1 | 0.0020 | 0.0004 | 0.22 |
| Glutathione peroxidase 3 OS=Homo sapiens OX=9606 GN=GPX3 PE=1 SV=2 | P22352 | GPX3 | 0.0005 | 0.0001 | 0.21 |
| Myosin light chain kinase, smooth muscle OS=Homo sapiens OX=9606 GN=MYLK PE=1 SV=4 | Q15746 | MYLK | 0.0029 | 0.0006 | 0.20 |
| Collagen alpha-2(IV) chain OS=Homo sapiens OX=9606 GN=COL4A2 PE=1 SV=4 | P08572 | COL4A2 | 0.0011 | 0.0002 | 0.20 |
| Latent-transforming growth factor beta-binding protein 4 OS=Homo sapiens OX=9606 GN=LTBP4 | Q8N2S1 | LTBP4 | 0.0001 | 0.0000 | 0.18 |
| Follistatin-related protein 1 OS=Homo sapiens OX=9606 GN=FSTL1 PE=1 SV=1 | Q12841 | FSTL1 | 0.0003 | 0.0000 | 0.13 |
| Keratin, type I cytoskeletal 9 OS=Homo sapiens OX=9606 GN=KRT9 PE=1 SV=3 | P35527 | KRT9 | 0.0007 | 0.0001 | 0.12 |
| Isoform 7 of Tropomyosin alpha-1 chain OS=Homo sapiens OX=9606 GN=TPM1 | P09493 | TPM1 | 0.0244 | 0.0025 | 0.10 |
| Tensin-1 OS=Homo sapiens OX=9606 GN=TNS1 PE=1 SV=2 | Q9HBL0 | TNS1 | 0.0004 | 0.0000 | 0.07 |
| Histone H1.0 OS=Homo sapiens OX=9606 GN=H1F0 PE=1 SV=3 | P07305 | H1F0 | 0.0060 | 0.0002 | 0.04 |
| Actin, aortic smooth muscle OS=Homo sapiens OX=9606 GN=ACTA2 PE=1 SV=1 | P62736 | ACTA2 | 0.0287 | 0.0011 | 0.04 |
| Prostatic acid phosphatase OS=Homo sapiens OX=9606 GN=ACPP PE=1 SV=3 | P15309 | ACPP | 0.0061 | 0.0002 | 0.03 |

**Table S3:** List of proteins observed to be overexpressed using urine proteomics by at least two folds in urine samples of patients with prostate cancer

| **Protein** | **Accession No.** | **Gene code** | **Control (Normalized and averaged)** | **Sample (Normalized and averaged)** | **Fold change in expression (Sample/Control)** |
| --- | --- | --- | --- | --- | --- |
| Lymphocyte antigen 6D OS=Homo sapiens OX=9606 GN=LY6D PE=1 SV=1 | Q14210 | LY6D | 0.00002 | 0.00032 | 17.47 |
| Cell surface glycoprotein MUC18 OS=Homo sapiens OX=9606 GN=MCAM PE=1 SV=2 | P43121 | MCAM | 0.00004 | 0.00021 | 4.64 |
| Isoform 4 of Latent-transforming growth factor beta-binding protein 1 OS=Homo sapiens OX=9606 GN=LTBP1 | Q14766 | LTBP1 | 0.00008 | 0.00029 | 3.86 |
| Isoform 2 of Transmembrane protease serine 2 OS=Homo sapiens OX=9606 GN=TMPRSS2 | O15393 | TMPRSS2 | 0.00007 | 0.00021 | 3.21 |
| Pigment epithelium-derived factor OS=Homo sapiens OX=9606 GN=SERPINF1 PE=1 | P36955 | SERPINF1 | 0.00004 | 0.00012 | 3.18 |
| Ras-related protein Rab-1A OS=Homo sapiens OX=9606 GN=RAB1A PE=1 SV=3 | P62820 | RAB1A | 0.00011 | 0.00023 | 2.05 |
| Interleukin-10 receptor subunit beta OS=Homo sapiens OX=9606 GN=IL10RB PE=1 SV=2 | Q08334 | IL10RB | 0.00007 | 0.00015 | 2.03 |

**Table S4:** List of proteins observed to be underexpressed using urine proteomics by at least two folds in urine samples of patients with prostate cancer

| **Protein** | **Accession No.** | **Gene code** | **Control (Normalized and averaged)** | **Sample (Normalized and averaged)** | **Fold change in expression (Sample/Control)** |
| --- | --- | --- | --- | --- | --- |
| Carboxypeptidase Z OS=Homo sapiens OX=9606 GN=CPZ PE=1 SV=2 | Q66K79 | CPZ | 0.00015 | 0.00008 | 0.51 |
| CD99 antigen-like protein 2 OS=Homo sapiens OX=9606 GN=CD99L2 PE=1 SV=1 | Q8TCZ2 | CD99L2 | 0.00119 | 0.00053 | 0.44 |
| Oxidized low-density lipoprotein receptor 1 OS=Homo sapiens OX=9606 GN=OLR1 PE=1 SV=1 | P78380 | OLR1 | 0.00051 | 0.00022 | 0.44 |
| Serine protease inhibitor Kazal-type 1 OS=Homo sapiens OX=9606 GN=SPINK1 PE=1 SV=2 | P00995 | SPINK1 | 0.00011 | 0.00005 | 0.43 |
| Isoform 4 of Methanethiol oxidase OS=Homo sapiens OX=9606 GN=SELENBP1 | Q13228 | SELENBP1 | 0.00012 | 0.00005 | 0.41 |
| Vitronectin OS=Homo sapiens OX=9606 GN=VTN | P04004 | VTN | 0.00386 | 0.00159 | 0.41 |
| Isoform 2 of Annexin A2 OS=Homo sapiens OX=9606 GN=ANXA2 | P07355 | ANXA2 | 0.00006 | 0.00002 | 0.40 |
| Isoform 5 of Acyl-CoA-binding protein OS=Homo sapiens OX=9606 GN=DBI | P07108 | DBI | 0.00033 | 0.00013 | 0.39 |
| Retinoid-inducible serine carboxypeptidase OS=Homo sapiens OX=9606 GN=SCPEP1 PE=1 SV=1 | Q9HB40 | SCPEP1 | 0.00032 | 0.00012 | 0.39 |
| Alpha-galactosidase A OS=Homo sapiens OX=9606 GN=GLA PE=1 SV=1 | P06280 | GLA | 0.00012 | 0.00004 | 0.35 |
| Thrombospondin-1 OS=Homo sapiens OX=9606 GN=THBS1 PE=1 SV=2 | P07996 | THBS1 | 0.00162 | 0.00036 | 0.22 |
| Podocalyxin OS=Homo sapiens OX=9606 GN=PODXL PE=1 SV=2 | O00592 | PODXL | 0.00024 | 0.00005 | 0.22 |
| Filamin-A OS=Homo sapiens OX=9606 GN=FLNA PE=1 SV=4 | P21333 | FLNA | 0.00033 | 0.00007 | 0.20 |
| Latent-transforming growth factor beta-binding protein 4 OS=Homo sapiens OX=9606 GN=LTBP4 PE=1 SV=2 | Q8N2S1 | LTBP4 | 0.00097 | 0.00019 | 0.19 |
| Protein delta homolog 2 OS=Homo sapiens OX=9606 GN=DLK2 PE=2 SV=1 | Q6UY11 | DLK2 | 0.00009 | 0.00001 | 0.14 |
